# Chromatin compaction and spatial organization in rice interphase nuclei

**DOI:** 10.1101/2023.06.28.546909

**Authors:** Alžběta Doležalová, Denisa Beránková, Veronika Koláčková, Eva Hřibová

**Affiliations:** Institute of Experimental Botany of the Czech Academy of Sciences, Centre of Plant Structural and Functional Genomics, 77900 Olomouc, Czech Republic

**Author notes:** Corresponding Author: Eva Hřibová Institute of Experimental Botany Šlechtitelů 31 779 00 Olomouc Czech Republic. E-mail addresses: Doležalová A. - Beránková D. - Koláčková V.

**Keywords:** 3D Immuno-FISH, chromosome painting, chromosome territory, rice, spatial organization, microscopy

## Abstract

Chromatin organization and its interactions are essential for biological processes such as DNA repair, transcription, and DNA replication. Detailed cytogenetics data on chromatin conformation, and the arrangement and mutual positioning of chromosome territories in interphase nuclei are still widely missing in plants. In this study, level of chromatin condensation in interphase nuclei of rice (*Oryza sativa*) and the distribution of chromosome territories (CTs) were analyzed. Super-resolution, stimulated emission depletion (STED), microscopy showed different level of chromatin condensation in leaf and root interphase nuclei. 3D immuno-FISH experiments with painting probes specific to chromosomes 9 and 2 were conducted to investigate their spatial distribution in root and leaf nuclei. Six different configurations of chromosome territories, including their complete association, weak association, and complete separation, were observed in root meristematic nuclei, and four configurations were observed in leaf nuclei. The volume of CTs and frequency of their association varied between the tissue types. The frequency of association of CTs specific to chromosome 9, containing NOR region, is also affected by the activity of the 45S rDNA locus. Our data suggested that the arrangement of chromosomes in the nucleus is connected to the position and the size of the nucleolus.

**Highlight:** Super-resolution STED microscopy uncovered detailed chromatin ultrastructure; high level of differences in chromatin condensation and mutual positioning of chromosome territories between and within leaf and root meristem G1 were observed.

## Introduction

Nuclear DNA is condensed together with structural proteins into higher-order chromatin structures, which serve as substrates for important biological processes such as DNA replication, transcription, and genome repair (Misteli, 2020). While the chromatin is packed into visible, highly condensed chromosome structures during mitosis, it is decondensed in the interphase of the cell cycle, and the borders of individual chromosomes can not be recognized. The fundamental questions are: how is the chromatin packed into chromosomes, how are the chromosomes organized during the interphase of the cell cycle, and how does the chromatin packing and chromosome positioning influence biological processes?

The organization of chromatin during the interphase can be analyzed by two methodological approaches: by high-throughput chromosome conformation capture (Hi-C) technique, followed by polymer modeling (Lieberman-Aiden *et al*., 2009; Giorgetti *et al*., 2014; Gibcus *et al*., 2018), and by three-dimensional fluorescence in situ hybridization (3D-FISH) and microscopic techniques (Bass *et al*., 1997). The Hi-C method combines 3C technique (Dekker *et al*., 2002) and next-generation sequencing (Lieberman-Aiden *et al*., 2009) to find out chromatin compaction. Recently, Hi-C techniques have been used in many living organisms to describe chromosome contact patterns, genome packing, and 3D chromatin architecture at much higher resolution (tens to hundreds of kilobases) than is provided by 3D-FISH (Dong *et al*., 2018; Dumur *et al*., 2019; Concia *et al*., 2020; Golicz *et al*., 2020). On the other hand, majority of Hi-C studies in plant species were performed on pooled tissues and thus could not provide information about the variability in spatial organization of individual chromosomes in 3D space of the interphase nuclei (Wang *et al*., 2015; Dong *et al*., 2017; Concia *et al*., 2020). This information can be achieved by application of recently developed cytogenetic techniques, oligo-painting, and 3D-FISH, which enable to visualize individual genome regions in 3D space of nuclei (Howe *et al*., 2013; Han *et al*., 2015).

Hi-C studies in metazoans and mammals revealed the existence of megabase-long chromatin compartments containing either active and open chromatin (A compartments), or inactive and closed chromatin (B compartments). Hi-C also allowed to describe organization into smaller (in average 800 kb long in mammals), self-interacting topologically associated domains (TADs), regulatory landscapes of chromosomes, which were revealed in animal interphase nuclei (e.g. Sexton and Cavalli, 2015; Ramírez *et al*., 2018; Szabo *et al*., 2018). Genes belonging to the same TADs display similar expression dynamics suggesting that their physical association is functionally related to gene expression control (de Graaf et al., 2013). In plants, 3D chromatin architecture is different. For instance, TADs were not observed in *A. thaliana* (Feng *et al*., 2014; Wang *et al*., 2015; Liu *et al*., 2017), instead their presence seems to be linked to species with larger genomes (Dong *et al*., 2017; Liu *et al*., 2017; Concia *et al*., 2020; Golicz *et al*., 2020). Since TADs have not been recognized in all plant species, the question arises whether they play an important role in the dynamics of plant chromosomes.

Complementary cytogenetic data to Hi-C studies are still missing. In plant research, chromosome distribution in interphase nuclei was studied by FISH with probes specific to functional chromosome domains such as centromeres and telomeres (Hou *et al*., 2018; Liu *et al*., 2020). The aim of these studies was to confirm the first microscopic observations *done by* Carl Rabl, who predicted that chromosome positioning in interphase nuclei follows their orientation in the preceding mitosis (Rabl, 1885; reviewed by Cremer *et al*., 2006). The so-called Rabl configuration, with centromeres and telomeres oriented on opposite poles of nuclei, was originally assigned to plants with large genomes (wheat, rye, barley) (Cremer *et al*., 2001). The concept of a Rabl-like pattern in plant species with large genomes and non-Rabl organization of interphase chromosomes in plants with small and medium genomes has been disproved early after it was proposed (Fujimoto *et al*., 2005). In rice, the majority of nuclei in somatic cells lack Rabl configuration (Prieto *et al*., 2004; Santos and Show, 2004; Němečková *et al*., 2020), however, chromosomes of pre-meiotic cells in anthers or xylem-vessel precursor cells seem to assume the Rabl configuration (Prieto *et al*., 2004; Santos and Show, 2004). Compared to numerous studies on the centromere-telomere organization in plant interphase nuclei (Fujimoto *et al*., 2005; Idziak *et al*., 2015; Nemečková *et al*., 2020; Shan *et al*., 2021), the visualization of the spatial positioning of individual chromosomes during interphase stays widely unknown. The mutual position of chromosomes during interphase was studied in *Arabidopsis thaliana* using BAC pools-based chromosome painting technique, showing that individual chromosomes tend to occupy separated territories (Pečinka *et al*., 2004). The extremely small genome of Arabidopsis is characterized by a specific, rosette-like, chromosome configuration (Armstrong *et al*.,2001; Fransz *et al*., 2002), which was not observed in any other plant species, thus we can not expect that the chromosome organization and dynamics revealed in *Arabidopsis* is universal to other plant species. Robaszkiewicz *et al*. (2016) later analyzed chromosome positioning in the 3D space of *Brachypodium distachyon*, which possesses Rabl orientation, and provided the first insight into the large variability of the interphase chromosome organization. However, the high level of variability in mutual chromosome organization shown in the study, could have been caused by the use of nuclei isolated from the pooled root tissue (Robaszkiewicz *et al*., 2016).

Our present study provides the first insight into chromatin compaction and variability of the spatial organization of CTs during the interphase of the cell cycle in highly dynamic root meristematic cells and diversified leaf nuclei. The use of super-resolution STED microscopy revealed different levels of chromatin compaction in root and leaf nuclei. 3D immuno-FISH experiments with chromosome-specific painting probes showed different types of mutual CTs positioning which varied between the root and leaf interphase nuclei.

## Material and methods

### Plant material, seeds germination and sample preparation

Seeds of rice (*Oryza sativa*) cultivar Nipponbare (2n=2x=24) were obtained from Prof. Ohmido Nobuko, Kobe University, Japan. Seeds were soaked in distilled water and bubbled for 24 h. After that, seeds germinated in a biological incubator at 24°C in a glass Petri dish on moistened filter paper until the primary roots were 3-4 cm long. Suspension of intact nuclei was prepared according to Doležel *et al*. (1992). Briefly, root tips were cut and fixed with 2% (v/v) formaldehyde in Tris buffer (10 mM Tris, 10 mM Na2EDTA, 100 mM NaCl, 0.1% Triton X-100, 2 % formaldehyde, pH 7.5) at 4°C for 30 min and washed three times with Tris buffer at 4°C. Meristematic parts of root tips (∼1 mm long) were excised from 70 roots per sample. Root meristems were homogenized in 500 µl LB01 buffer (Doležel *et al*., 1989) by Polytron PT 1200 homogenizer (Kinematica AG, Littua, Switzerland) for 13 s at 14 500 rpm. Finally, the suspensions were filtered through a 20 µm nylon mesh and analyzed using a FASCAria II SORP flow cytometer and sorter (BD Bioscience, San Jose, USA). Nuclei representing the G1 phase of the cell cycle were sorted into 1x meiocyte buffer (Bass *et al.,* 1997; Howe *et al.,* 2013).

### Root microtome sectioning and FISH

Roots fixed with 2% (v/v) formaldehyde in Tris were embedded in Cryo-Gel (Leica Biosystems, ID:39475237) and cut onto a cryostat (Leica CM1950) at a thickness of 20 µm. The resulting segments were transferred to super-frost slide (Thermo Scientific). Slides were allowed to dry overnight at room temperature and then either immediately utilized for FISH or stored at 4 °C until use. Prior to FISH, slides with root segments in cryo-gel were washed for 10 min in 1x PBS and subsequently dehydrated in ethanol series (70%, 85%, 100% ethanol), each for 2 min. Hybridization mix (50 µl) containing 50% (v/v) formamide, 10% (w/v) dextran sulfate in 2x SSC, 1 µg sheared salmon sperm DNA (Invitrogen, AM9680) and 200 ng per probe was added onto the slides and denatured for 8 min at 78°C and cooled slowly (50 °C 1 min, 45 °C 1 min, 40 °C 1 min, 38 °C 5 min). After that, slides were hybridized overnight at 37 °C. The next day, slides were washed 3×5 min in 4xSSC, and root sections were counterstained with DAPI in VECTASHIELD Antifade Mounting Medium (Vector Laboratories, Burlingame, CA, USA).

### Probes for FISH

Oligonucleotides specific for individual chromosomes were identified in the reference genome sequence of *Oryza sativa* cv. Nipponbare (version_7.0; http://rice.uga.edu/; Kawahara et al., 2013) using the Chorus v2 program pipeline (Zhang *et al*., 2021). Two sets of oligomers were synthesized by Arbor Biosciences (Ann Arbor, Michigan, USA). Labeled oligomer probes were prepared according to Han *et al*. (2015). Probes specific for the long and short arms of chromosome 2 were labeled by biotin-16-dUTP and by aminoallyl-dUTP-CY3, and chromosome 9 was labeled by digoxigenin-11-dUPT and aminoallyl-dUTP-CY5 (Jena Biosciences, Jena, Germany). The painting probe of longer chromosome 2 contained 40,000 unique 45-mers and the painting probe specific to short chromosome 9 contained 20,000 unique 45-mers. Probes specific for 45S ribosomal DNA were amplified using specific primers (Ohmido and Fukui, 1995) and directly labeled with aminoallyl-dUTP-CY5 (Jena Biosciences, Jena, Germany).

### Immuno-staining and fluorescence in situ hybridization (FISH)

Flow sorted nuclei were mounted in polyacrylamide gel according to Němečková *et al*. (2020). To visualize 45S rDNA, chromosome 2 and chromosome 9 together with fibrillarin, staining procedures, and washes were performed according to Němečková *et al*. (2020). Primary antibody anti-fibrillarin was diluted at 1:100 (ab4566, Abcam, Cambridge, UK). The hybridization mix for FISH contained 400 ng of individual probes.

### Spirochrome staining and sample preparation for STED

Flow sorted nuclei were mounted in polyacrylamide gel onto silane-cover glass. High-precision cover glasses were prepared according to de Almeida Engler (2001) with modifications. Slides were washed in water for 15 min and in ethanol for 30 min. Slides were air dried for 10 minutes and then freshly prepared 2% 3-aminopropyltriethoxysilane (Sigma) in acetone was applied for 30 min. Slides were washed twice in distilled water, dried overnight at 37°C, and stored at room temperature (RT). After gel polymerization, polyacrylamide pads were washed in MBA buffer (Howe *et al*., 2013; Bass *et al*., 2014) and let dry at RT. Next, glycerol mounting medium AD-MOUNT S (ADVi, Říčany, Czech Republic) with SPY650-DNA (diluted 1:1000) (Spirochrome AG, cat#: SC501, Stein am Rhein, Switzerland) was applied onto the pads and covered with a microscopic slide.

### Confocal and STED microscopy, and image analysis

Images were acquired using Leica TCS SP8 STED 3X confocal microscope (Leica Microsystems, Wetzlar, Germany) equipped with 63x/1.4 NA Oil Plan Apochromat objective and Leica LAS-X software with Leica Lightning module. Image stacks were captured separately for each chromosome using 647 nm, 561 nm, 488, and 405 nm laser lines for excitation and appropriate emission filters. Typically, an image stack of about 50 slides with 0.15 µm spacing was acquired. Root sections were acquired via the Navigator module using a 63x objective and the final picture was created by the mosaic merge function. Different chromatin structure of leaf and root was captured in the STED mode with *100x 1.4 NA STED oil objective*. The pinhole was set to 0.75 AU. The resolution was estimated using LAS-X software according to full width at half maximum criterion. The chromatin signal labeled by spirochrom (SPY650-DNA) was captured with a lateral resolution of c. 52 nm. LAS-X software was also used to produce color heat maps of individual nuclei.

3D models of microscopic images and volume calculations were performed using Imaris 9.7 software (Bitplane, Oxford Instruments, Zurich, Switzerland). The volume of each nucleus, nucleolus, and chromosome territories was estimated based on the primary intensity of fluorescence obtained by microscopy. Imaris function ‘Surface’ was used for modeling the chromosome arrangement in the nucleus and for modeling the 45S rDNA, chromosomes, and fibrillarin. Chanel contrast was adjusted using ‘Chanel Adjustment’ and videos were created using the ‘Animation function’. About 100 nuclei were analyzed for each selected variant.

## Results

### Variation in chromatin condensation and nuclei features

To analyze and compare the level of chromatin condensation in G1 interphase nuclei of young leaves and root meristems, we applied stimulated emission depletion (STED) microscopy. With this aim, mildly fixed flow sorted G1 nuclei from leaves and root meristems were mounted in polyacrylamide gel onto silane-coated high-precision cover glass to ensure their 3D structure will be preserved. STED analysis uncovered detailed chromatin ultrastructure and revealed differences in the level of chromatin compaction between G1 nuclei isolated from leaves and root meristems. G1 nuclei of root meristem, which undergo repeated and rapid cell division, were characterized by more relaxed chromatin and apparent ultra-structures (Figure 1A). In comparison, a more compact structure of chromatin and presence of lower amount of interchromatin compartments was found in G1 nuclei isolated from differentiated leaf cells (Figure 1A). The chromatin condensation in G1 nuclei isolated from both tissues were also visualized as color heat maps (e.g. Cremer *et al.,* 2017; Cremer and Cremer, 2019), which display differences in the general chromatin organization between root meristem and leaf G1 nuclei (Figure 1A). The width of chromatin fiber in G1 nuclei from leaf reached 240 nm on average, while in root, the chromatin fiber was three times narrower, about 83 nm in width (Figure 1B)

**Figure 1.**
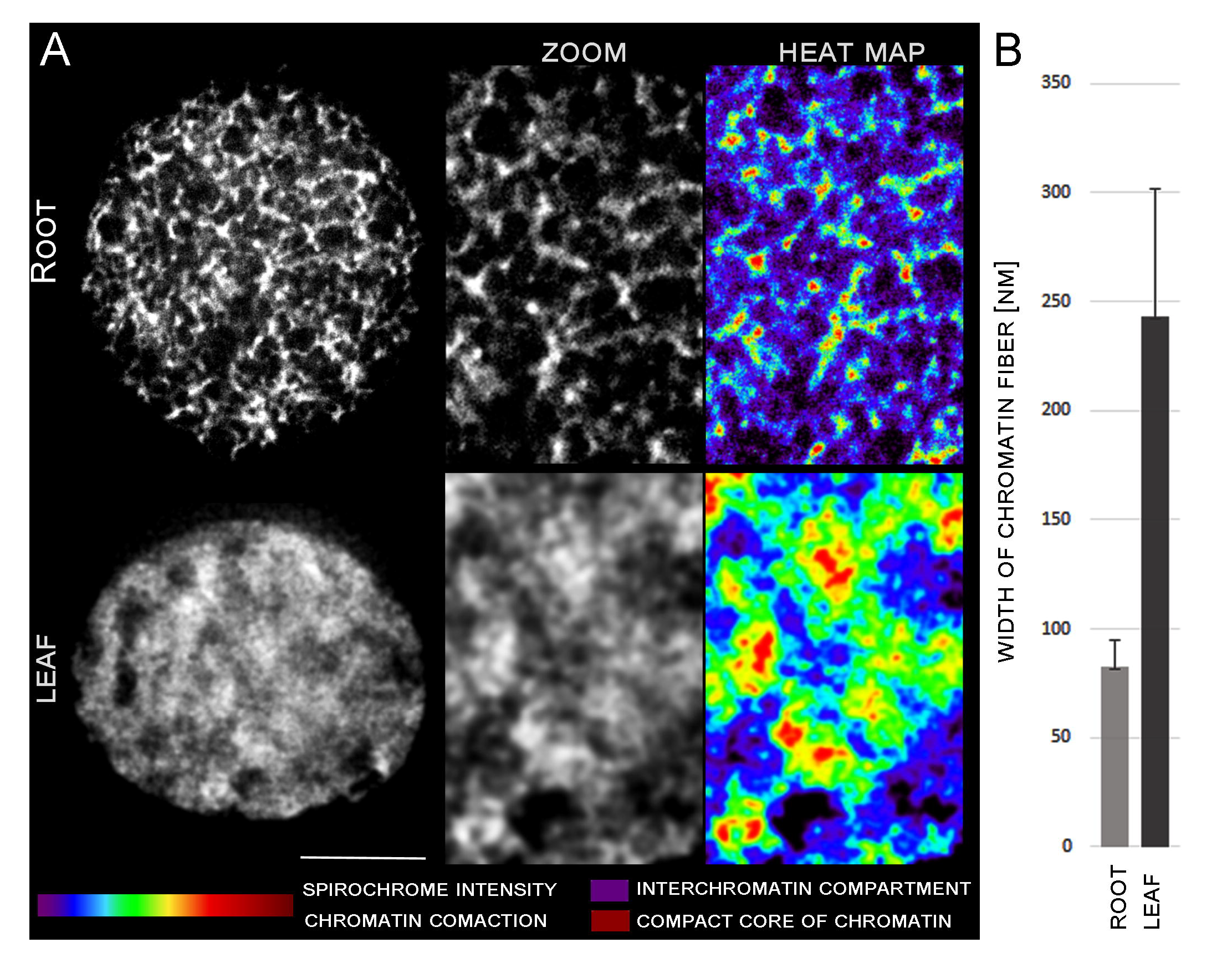
Chromatin condensation in G1 nuclei of young leaves and root meristem. DNA was stained by spirochrome (white). Differences in DNA structure are clearly visible in zoomed pictures and heat maps (A). (B) Graph of chromatin fiber width measurement. Bar 2 µm.

Likewise, the nuclei volume of G1 nuclei isolated from root meristematic zones was more than three times higher (199 µm3) compared to leaf nuclei (59.6 µm3) (Table 1). Similarly, volumes of nucleoli, which were visualized by immunodetection with nucleolus-specific protein fibrillarin, varied between leaves and root meristem. The volume of root nucleoli occupied 14.13 µm3 on average, which represents 7.1 % of the volume of the root nucleus (Supplementary video 1). Leaf nucleolus occupied 0.7 µm3, representing only 1.2 % of the leaf nucleus. (Table 1; Supplementary video 2).

Further analyses of about 200 G1 nuclei specific for both analyzed tissues revealed also variation in their shapes (Figure 2A). Majority of the nuclei had elliptical shape (∼ 67 %), and the rest of the G1 nuclei had spindle-like (∼ 21 %) and donut-like (∼ 13 %) shapes in root meristematic cells. Proportion of the G1 nuclei shapes was almost identical for both studied tissues (Table 1, Figure 2A).

**Figure 2.**
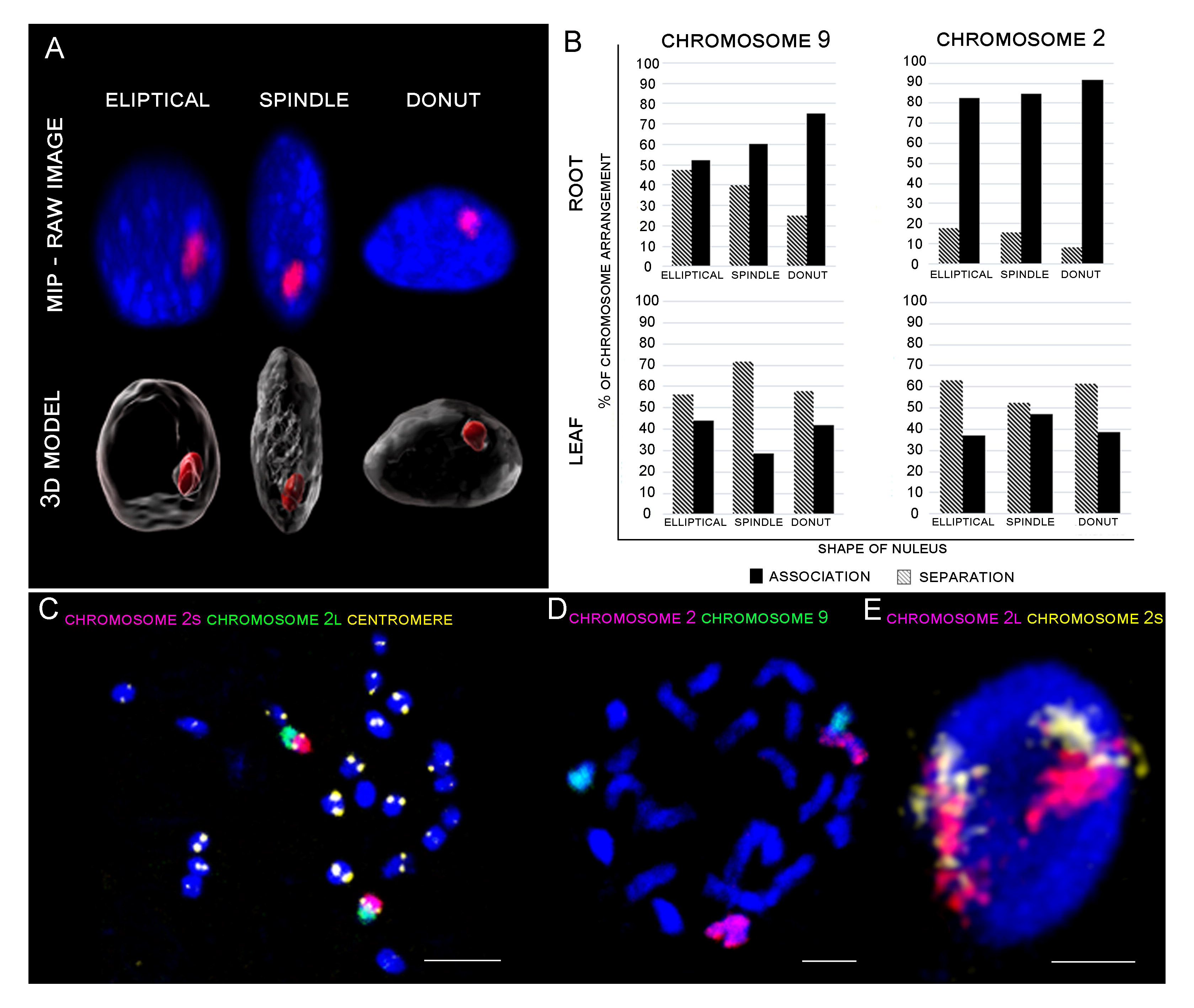
Representative figures of oligo-painting FISH and immunolabeling. (A) Differences in shape of analyzed nuclei (nuclear DNA stained by DAPI, blue). Nucleolus was visualized using fibrillarin immunolabeling (red). (B) Correlation between shape of the nucleus and CTs association. (C) Visualization of centromere (yellow), short arm of chromosome 2 (2S) (pink), and long arm (2L) (green) on metaphase chromosomes. (D) Visualization of chromosome 2 (pink) and chromosome 9 (green) by oligo-painting FISH on prometaphase chromosomes. Chromosomes were counterstained with DAPI (blue). (E) Maximal intensity projection of G1 nuclei. Two separate chromosome territories correspond to two homologues chromosomes. Long arm (2L) of chromosome in pink, short arm of chromosome (2S) in yellow. DNA was counterstained with DAPI (blue). Bar 3 µm.

### Chromosome specific painting probes

With the aim to analyze positioning of whole chromosomes during the interphase, we prepared oligo-painting probes for two rice chromosomes. Based on the previous Hi-C results, which proposed presence of two sets of chromosomes differing in level of their association (Dong *et al*., 2018), we analyzed detailed positioning of two chromosomes representing the two different sets. Long, sub-metacentric chromosome 2 (member of chromosome set which showed close association), and short acrocentric chromosome 9 containing NOR region and belonging to the set of chromosomes which did not show apparent association (Dong *et al*., 2018). Painting probes specific for both chromosomes were designed from a set of non-overlapping unique oligomers identified in reference genome sequence of *Oryza sativa* cv. Nipponbare v.7.0 (Kawahara *et al*., 2013) using Chorus v2 program pipeline (Zhang *et al*., 2021). Both oligos libraries were designed to achieve a density of at least 0.9 oligo per kb to ensure good visibility of hybridization signals after FISH. Sensitivity and suitability of the painting probe for chromosome identification *in situ* was confirmed by FISH on prometaphase and metaphase chromosomes (Figure 2C, 2D), and further on flow sorted G1 nuclei of root (Figure 3A, 3C, 3D) meristem and leaf tissue (Figure 3B). Both probes produced bright signals without any cross-hybridization. As expected, painting probes were strongly localized to chromosome arms, and centromeric regions and NOR region (containing specific repetitive DNA sequences) remained without good visible signal (Figure 2C, 2D).

**Figure 3.**
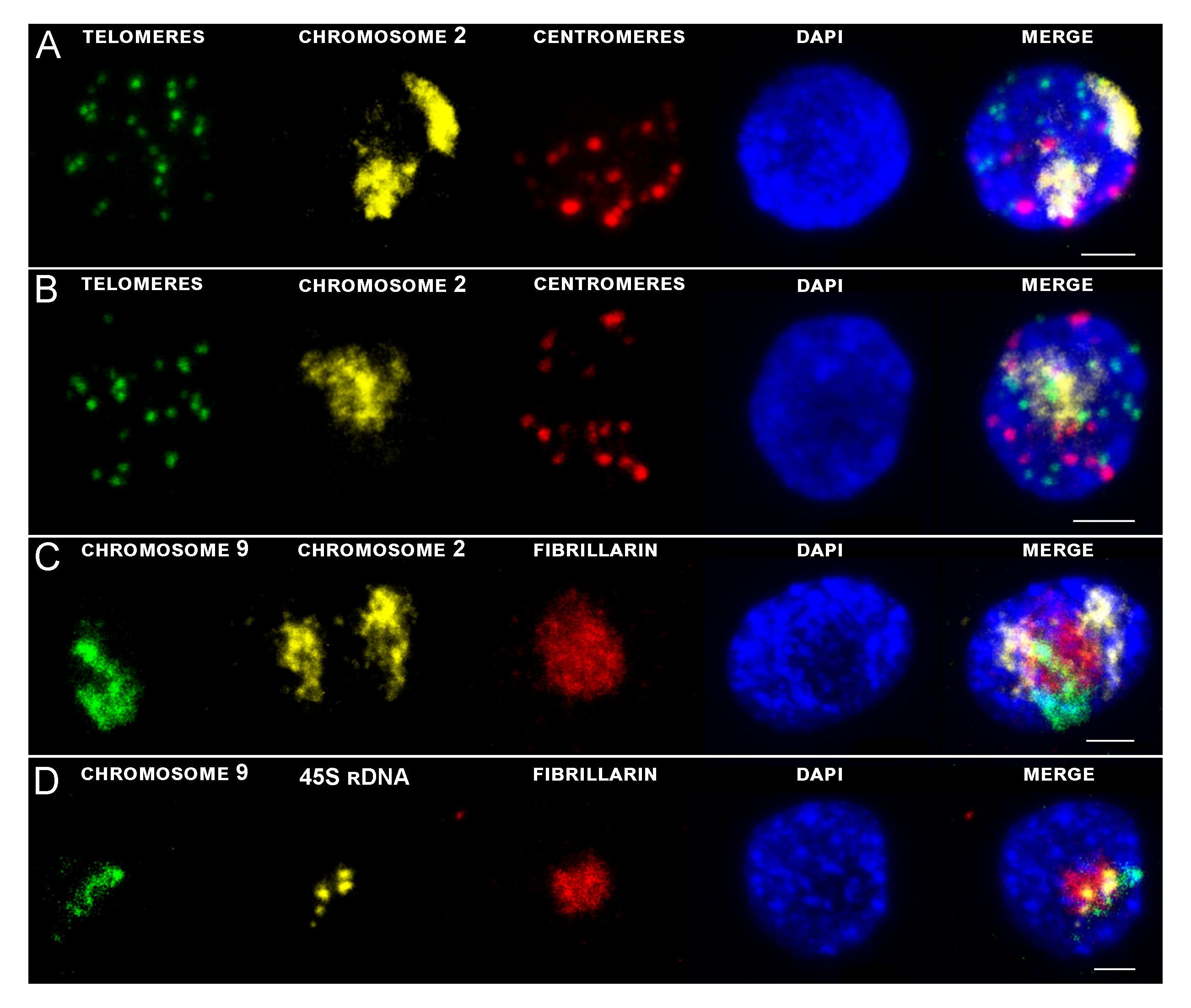
Maximal intensity projection of G1 nuclei of rice with immuno-FISH localization of different specific probes on flow sorted G1 nuclei of root meristem (A, C, D) and leaf tissue (B). DNA was counterstained with DAPI (blue). Bar 2 µm.

### Mutual position of chromosomes in G1 interphase nuclei

3D-FISH with the chromosome painting probes on G1 nuclei of rice revealed presence of compact structures in both examined tissue types and confirmed presence of so-called chromosome territories (CTs), which were predicted by Hi-C studies (Dong *et al*., 2018). Painting FISH revealed variability in constitution of the CTs, which were present either as two separated territories corresponding to two homologous chromosomes in G1 nuclei (Figure 2E, 3A, 3C), or as one large territory in which homologous chromosomes were tightly connected (Figure 3B). In general, higher proportion of G1 nuclei isolated from leave tissue showed close association of homologous chromosomes which were visualized as one large CT (63 % for chromosome 9; and 59 % for chromosome 2) compared to root meristem, which mostly contained G1 nuclei with two separated CTs (87 % for chromosome 9; and 59 % for chromosome 2) (Figure 4).

**Figure 4.**
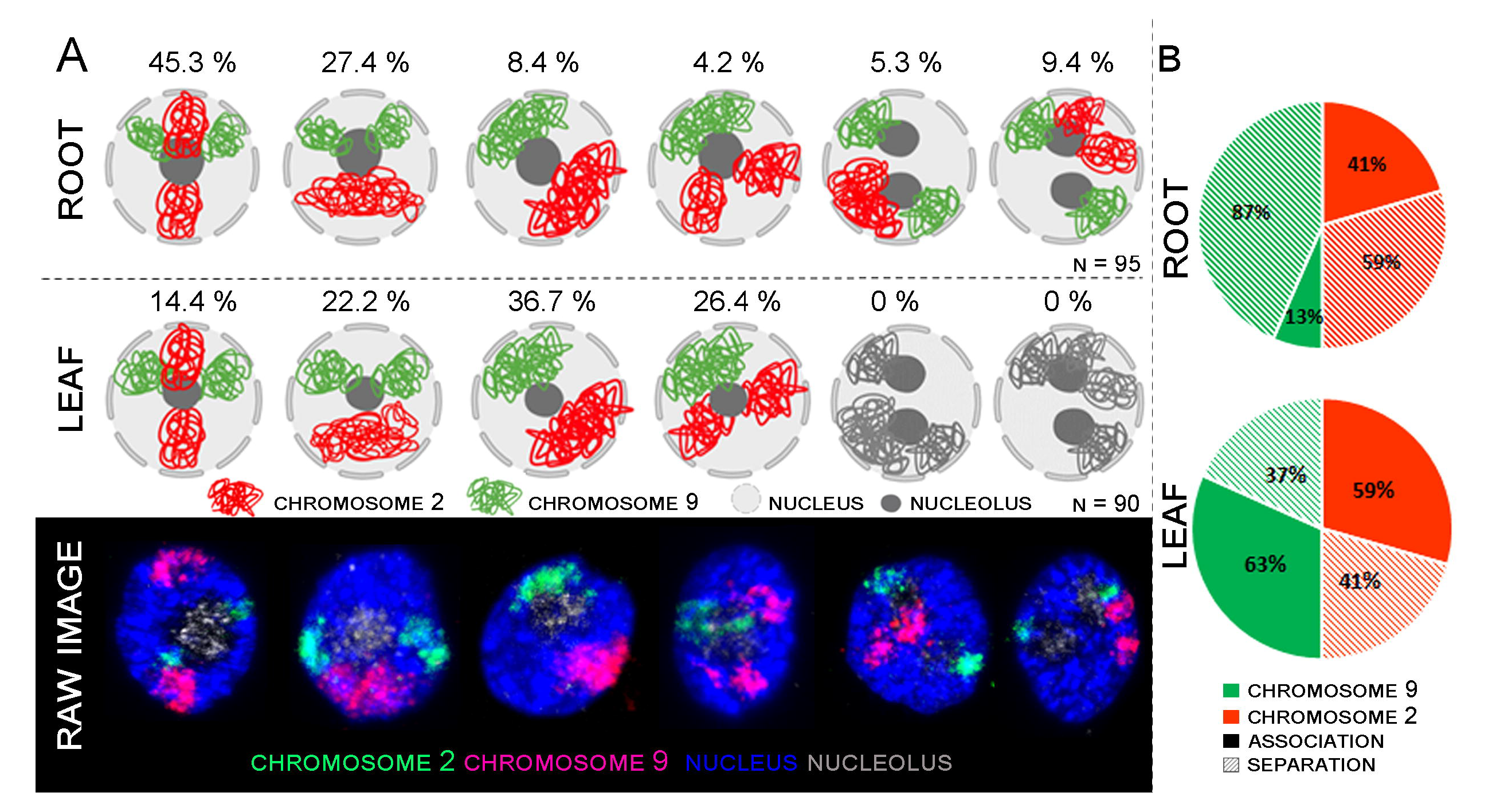
Comparison between root and leaf chromosome arrangement. (A) Models of individual arrangements created with BioRender.com. based on raw data observation. (B) Graph of both chromosome association comparison. Association and separation in displayed in root and leaf tissue.

Comparison of both territory volumes corresponding to homologous chromosomes did not reveal significant differences. Both chromosome territories represented 8.2 % of nucleus volume in root meristematic cells and 8.8% of the nucleus of leaf cells. The volume of separated territories of chromosome 2 was estimated to occupy approximately 4 % of the nucleus volume in both plant tissues (Table 2).

Co-hybridization and visualization of both chromosomes showed six different types of their mutual arrangement in G1 nuclei of root meristem (Figure 4A). About 45,3 % of examined root meristematic nuclei contained chromosomes 2 and 9 organized in two separate CTs concurrently, and additional 27,4 % of the nuclei contained chromosome 9 arranged in two separate CTs, and chromosome 2 in one large CT. 14,7 % of analyzed nuclei contained two nucleoli and in all these cases, CTs of chromosome 9 were separated. (Figure 4A).

In comparison, only four different arrangements of chromosome 2 and 9 were observed in leaf G1 nuclei. Nuclei containing two nucleoli were not present. 36.7 % of leaf nuclei contained two large CTs corresponding to chromosomes 2 and 9, other nuclei contained one large CT of chromosome 9 and two separated CTs specific to chromosome 2 (26.4 %). In the similar number of nuclei (22.2 %) CTs of homologous chromosome 2 were associated, and CTs of chromosome 9 were separated. Finally, 14.4 % of leaf G1 nuclei contained both chromosomes arranged in separate CTs (Figure 4A).

The difference in the proportion of separated and associated CTs between chromosomes 2 and 9 can be caused by the presence of NOR region on the short arm of chromosome 9. NOR region consists of 45S rRNA genes which constitute nucleoli, so the position of chromosome 9 in interphase nuclei also depends on the position and nature of the nucleolus/nucleoli (Supplementary Figure 1). A detailed 3D analysis revealed different numbers of 45S rDNA loci in the root and leaf. In the root, two major loci were usually observed on the periphery of the nucleolus and 2-4 small signals were observed inside the nucleolus (Supplementary Figure 1, Table 3). In comparison, only 1-2 45S rDNA loci situated on the periphery of the nucleolus were observed in the leaf, where the nucleolus occupies much lower volume (Supplementary Figure 1, Table 3). Detailed image analysis of leaf G1 nuclei revealed that the position of chromosome 2 is more random compared to chromosome 9. Large sub-metacentric chromosome 2 was in most cases arranged through the entire nucleus volume in the z-axis, with a large region located on the nuclear periphery (Figure 5). Even though, chromosome 2 does not contain rRNA genes and is not directly connected to nucleoli, its spatial positioning seems to be influenced by the nature of nucleoli (size, number, and position inside the nucleus).

**Figure 5.**
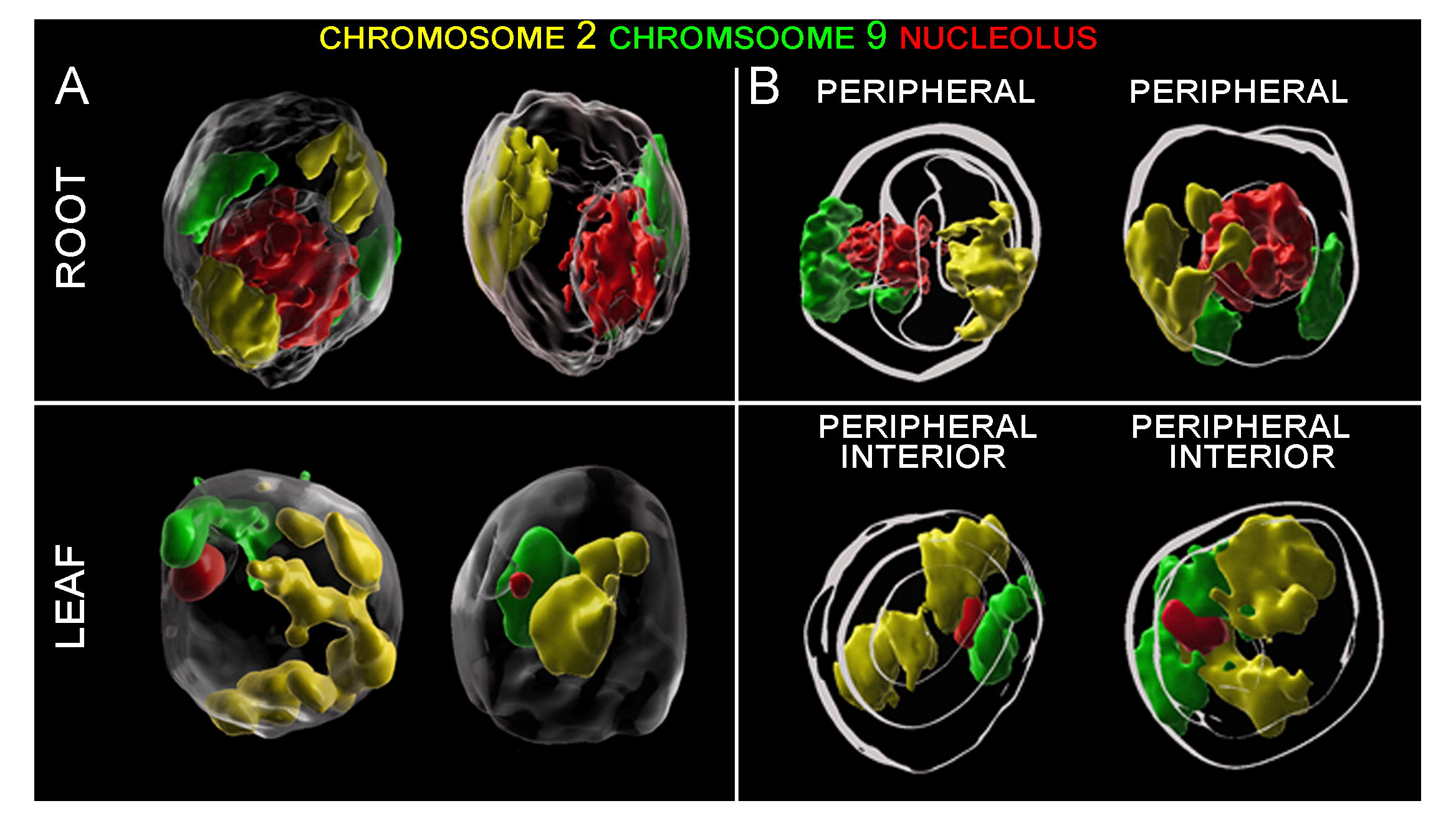
3D models of CTs positioning in root and leave G1 nuclei. (A) Spatial positioning of CTs specific to chromosome 2 (yellow) and 9 (green) and nucleoli (red). (B) Model showing spatial arrangement of the CTs and nucleoli with respect to the center and periphery of the nucleus. Shells of equal area depict regions of the nuclei, where signals of DAPI (white) and chromosome 2 (yellow) and chromosome 9 (green) were localized.

As we mentioned above, the G1 nuclei of both tissues varied also in their shape (Table 1, Figure 2A), thus we investigated relations between the CTs arrangement and nuclei shapes. Our data showed, that specific arrangements of CTs did not correlate with different shapes of nuclei. Despite lower proportion of spindle and donut-like nuclei (Table 1, Figure 2B), all CTs rearrangements (two separated territories, and homologous chromosome associated territory specific to chromosomes 2 and 9) were present in all examined nuclei (Figure 2B). Nevertheless, the potential connection between dominant pattern of CTs and the nuclear shape needs to be investigated in more detail on larger sample set due to unequal representation of spindle and donut-like nuclei (Table 1, Figure 2B).

Finally, we investigated the patterns of chromosome positioning in different cell types of the root meristem tissue. As we showed earlier, rice root meristematic cells did not show Rabl configuration of chromosomes during the interphase of the cell cycle (Němečková *et al*., 2020). The only exception was described by Prieto *et al*. (2004). They showed that xylem vessel cells, which are bigger and probably containing endoreduplicated nuclei, tend to achieve Rabl configuration. To confirm these findings by 3D FISH, we localized probes specific to centromeric and telomeric sequences on ultra-thin root sections prepared by cryomicrotome. Rabl configuration was observed in rice xylem vessel cells as well as in cortex cells (Figure 6A, B). Both cell types are bigger and the volume measurements of their nuclei performed in Imaris software indicate the presence of the endoreduplication. Unfortunately, proportion of these specific cell types in roots of rice is very low, so it is not possible to identify them by flow cytometry, estimate their DNA content and thus confirm presence of endoreduplication.

**Figure 6.**
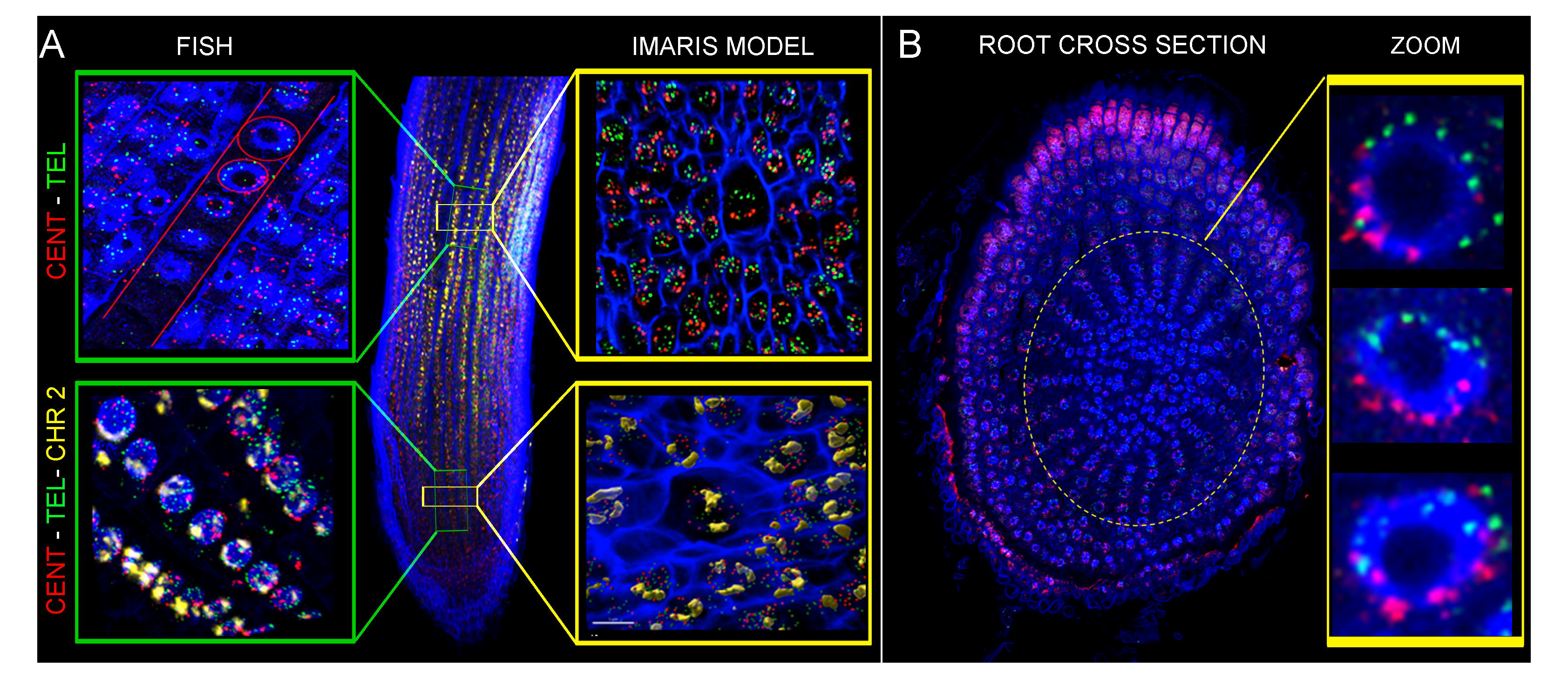
Oligo-painting FISH on root ultra-thin sections prepared by cryomicrotome. Centromeric probe (red), telomeric probe (green) and specific probe for chromosome 2 (yellow) were applied. Pictures displayed evidence of Rabl configuration in xylem (A) and in cortex (B) cells.

## Disscusion

Early studies of chromatin structure which used electron microscopy suggested its helical arrangement into 30 nm nucleosome fiber (Finch and Klug, 1976; Woodcock *et al*., 1984; Bordas *et al*., 1986). However, this model of chromatin folding and its higher-order organization became controversial due to the difference in observation between in vivo and in vitro conditions (Maeshima *et al*., 2019; Prieto et Maeshima, 2019). Recent development of super-resolution microscopy techniques, which enable to reach a resolution of about 1-250 nm (reviewed in Valli *et al*., 2021), allowed to describe a presence of 100-200 nm higher-order chromonema fibers (Kireeva *et al*., 2004; Maison *et al*., 2010; Belmont, 2014). Studies of DNA replication foci in human cells proposed a globular folding of chromatin with a diameter of about 110-150 nm (Jackson and Pombo, 1998; Albiez *et al*., 2006; Cseresnyes *et al*., 2009; Markaki *et al*., 2012).

In our study, we have analyzed chromatin compaction in the interphase nuclei of highly dynamic root meristematic cells and nuclei isolated from differentiated leaf cells. We have used STED super-resolution microscopy which can reach xy-resolution less than 60 nm, and also enables acquisition of three-dimensional images (Dumur *et al*., 2019; Moors *et al*., 2021; Frolikova *et al*., 2023). To provide information on chromatin compaction during the interphase of the cell cycle, mild formaldehyde fixation of the nuclei and their further mounting in polyacrylamide gel was used to preserve 3D chromatin structure and to avoid chromatin destruction during sample preparation (Bass *et al*., 2014; Howe *et al*., 2013; Němečková *et al*., 2020). Another important feature of the sample preparation for STED microscopy was the selection of appropriate mounting media, which would not have negative effect on 3D structure of the nuclei (Koláčková *et al*., in preparation).

Striking difference in chromatin compaction in G1 nuclei of root and leaf tissues were observed (Figure 1). Presence of 80 nm chromatin fibers was revealed in rice root meristematic G1 nuclei. Similar width of chromatin fiber (70 nm) was observed in metaphase chromosomes of Drosophila (Matsuda *et al*., 2010) and recently in mitotic chromosomes of barley root meristem (Kubalová *et al*., 2023). These results could indicate that the higher level of chromatin spiralization, which is typical for mitotic chromosomes, is maintained in interphase nuclei of highly dynamic meristematic cells. On the contrary, the diameter of rice leaf chromatin fibers was three times higher, reaching 240 nm. Similar variability in chromatin fibers was observed in human and animal studies, including metazoans (reviewed by Hansen et al., 2018). Actually, studies of Belmont *et al*. (1994) and Dehghani *et al*. (2005) showed presence of two classes of chromatin fibers, with diameters 60-80 nm, and 100-130 nm in early G1 and late G1/early S. Described diameter of rice higher order chromatin structure correlates with the diameter of the He-la cells’ higher-order chromatin structure (220 nm) (Nozaki *et al*., 2017). Root meristem and leaf nuclei varied also in the volume and level of chromatin compactness. Root meristem G1 nuclei were more than three times larger and consist of more relaxed chromatin with higher proportion of interchromatin compartments (Figure 1). As we analyzed G1 nuclei of highly dynamic root meristem cells, we can speculate, that the bigger size of these nuclei and higher proportion of interchromatin compartments are needed for the synthesis of mRNA and proteins, which are required for DNA synthesis in the following S phase. In comparison, differentiated cells of leaf tissues contained smaller G1 nuclei consisting of more compact chromatin with lower proportion of interchromatin compartments, where transcription takes a place, as was demonstrated in human studies (Hübner *et al*., 2015; Cremer et Cremer, 2019).

Recently, it was shown that nuclear architecture, the size and shape, and positioning of CTs during interphase, can be influenced by several factors, especially the size of a given chromosome, position of centromere, and the shape of nucleolus. In *Brachypodium distachyon*, a plant species that maintains Rabl configuration, a high level of homologous CTs associations was found in spherical nuclei, while it was negatively correlated with elongated nuclei (Robaszikewicz *et al*., 2016). Similar results were described for plants with rosette-like chromosome conformation in the nuclei of both, root and leaf, tissues (Pečinka *et al*., 2004). In our study, mutual position of two morphologically different chromosomes in interphase nuclei was not correlated with the nuclei shape. On the other hand, we revealed differences in organization and mutual chromosome position between root meristem and leaf G1 nuclei. We observed presence of discrete chromosome territories specific to both visualized chromosomes. CTs of NOR bearing chromosome 9 were mostly separated in root meristem nuclei, while their (CTs) association prevailed in leaf G1 nuclei, regardless their shape. Our findings are not in agreement with the organization and distribution of CTs of NOR bearing chromosome in *Brachypodium*, which were predominantly associated (59,3 %) (Robaszkiewicz *et al*., 2016). Robaszkiewicz *et al*. (2016) also suggested, that the length of a particular chromosome may influence the dominant pattern of its spatial arrangement inside the nucleus, and showed that CTs of the longest chromosome were usually associated. However, the high level of variability in mutual chromosome organization shown in the study of Robaszkiewicz *et al*. (2016) could be caused by the analysis of nuclei isolated from the pooled root tissue. Random positioning of most CTs was observed in *Arabidopsis*. The only exception was the position of NOR bearing chromosomes, which seemed to be connected to the position of nucleoli (Lysak *et al*., 2001; Pecinka *et al*., 2004; Berr and Schubert, 2007). Spatial organization and mutual position of CTs in 3D space of large plant genomes with Rabl configuration have not yet been analyzed by in situ techniques. The only exception was the visualization of alien chromosomes in wheat-rye and wheat-barley introgression lines (Koláčková *et al*., 2019; Perníčková *et al*., 2019). In both cases, a complete separation of CTs corresponding to alien chromosomes was observed in majority (83 – 89 %) of studied root meristem cells (Koláčková *et al*., 2019; Perníčková *et al*., 2019).

The discrepancies in CTs organization and positioning in 3D nuclear space between our work and previous studies, especially those of Robaszkiewicz *et al*. (2016), could be also caused by the difference of chromosome configuration (Rabl and non-Rabl) in the studied species. Further investigation has to be done to find out if chromosome configuration affects the organization and mutual position of CTs during the interphase of the cell cycle.

As we already mentioned, the shape and number of nucleoli represent another factor, which can affect the CTs positioning. Derenzini *et al*. (1998) showed, that cancer dividing cells produced elevated amounts of rRNA and often possessed large nucleoli whereas down-regulation of rRNA gene transcription led to reduction in nucleolar size. More recently, Tiku *et al*. (2018) showed, that size of the nucleolus positively correlates with rRNA synthesis. Analysis of purified nucleoli of *A. thaliana* showed that active rRNA genes are present within nucleoli whereas silent copies are excluded (Pontviane *et al*., 2013). Correlation between rRNA activity and arrangement of chromosome territories was indicated also in our study. Homologs of chromosome 9 were organized into separated territories (in 93 % of all events) in G1 nuclei of root meristem, where the rRNA genes are being highly expressed (Tulpová *et al*., 2022). On the other hand, chromosome 9 was more associated (59 % associated, 41 % separated) in leaf tissue, in which smaller volume of nucleolus and only 1-2 clusters of 45s rDNA were observed.

Our study showed high rate of variability in mutual chromosome positioning in the 3D space of G1 nuclei isolated from both plant tissues. This variability in the association/separation of homologous CTs may reflect the interphase chromatin dynamics. Movement of chromatin was described in *Arabidopsis* interphase nuclei by visualization of tagged loci in live seedlings (Kato and Lam, 2003), and in yeast (e.g. Heun, 2001; Bystricky *et al*., 2004; Hajjoul *et al*., 2013), animal and human cells (e.g. Chubb *et al*., 2002; Levi *et al*., 2005, Germier *et al*., 2017; Nozaki *et al*., 2023).

The observed heterogeneity in chromosome positioning and variability in chromatin condensation within different tissues explain the discrepancy between contact frequencies and distance distributions obtained by Hi-C and 3D-FISH (Fudenberg *et al*., 2017). In plants, most Hi-C studies, which can be also used to create putative models of chromatin condensation and chromosome positioning, were done on pooled tissues (Wang *et al*., 2015; Dong *et al*., 2017; Concia *et al*., 2020). Therefore, 3D modeling was performed based on averages of large numbers of cells, and the information on potential variability in 3D structure among different cells or cell types was lost. This can be overcome by single-cell Hi-C (scHi-C) experiments (Nagano *et al*., 2013; Ramani *et al*., 2017; Tan *et al*., 2018). In plant research, scHi-C experiments are not numerous. For instance, in rice, this technique was used to study variability in chromatin organization in eggs, sperm cells, unicellular zygotes, and shoot mesophyll cells. Even though the analysis was performed only on four cells representing each tissue type, theoretical models of chromosome folding and their mutual organization indicated variability in the positioning of chromosome territories among the analyzed nuclei (Zhou *et al*., 2019).

To conclude our study, we showed that advanced microscopy combined with recent cytogenetics techniques is a powerful tool for analysis and comparison of mutual chromosomes positions in the nuclei during the interphase of the cell cycle. Our experiments support the hypothesis, that chromatin organization is not determined by the shape of the nucleus. On the other hand, it appears that the size of the nucleolus and its position in the nucleus plays a role in chromosome positioning during interphase. The analysis of large number of nuclei confirms variability in chromosome organization into nuclear territories and their mutual positioning within and also between nuclei isolated from different tissue types.

Furthermore, the use of super-resolution STED microscopy corroborates striking differences in chromatin folding and organization in the interphase nuclei isolated from the two studied plant tissues.

## Abbreviations

3D: three-dimensional
CenH3: centromere-specific variant of histone H3
CT: chromosome territory
FISH: fluorescence *in situ* hybridization
HiC: high-throughput chromosome conformation capture
NOR: nucleolar organizer region
PAA: polyacrylamide
rDNA: ribosomal deoxyribonucleic acid
ROI: region of interest
RT: room temperature
STED: stimulated emission depletion
TADs: topologically associated domains

## Acknowledgments

We would like to cordially thank Jitka Weiserová and Dr. Petr Cápal for flow sorting, Zdeňka Dubská for excellent technical assistance, and Lucie Kobrlová for enabling the sample preparation on cryomicrotome and for the technical support. We would like to thank Dr. Jana Čížková for her valuable comments. The computing was supported by the project "e-Infrastruktura CZ" (e-INFRA CZ LM2018140) supported by the Ministry of Education, Youth and Sports of the Czech Republic, and by the ELIXIR-CZ project (LM2018131), part of the international ELIXIR infrastructure. This research was funded by the ERDF project "Plants as a tool for sustainable global development" (No. CZ.02.1.01/0.0/0.0/16_019/0000827).

## Author Contributions

E.H., and A.D. conceived the project. E.H. and D.B. designed and prepared painting probes, A.D., D.B., and V.K. conducted cytogenetic part of the work. A.D. and E.H., wrote the original draft, D.B. and V.K., reviewed and edited the manuscript. All authors have read and approved the final manuscript.

---

**Table 1** Analysis of all tested G1 nuclei specific for both analyzed tissues revealing variation in shape and volume of nucleus and nucleolus.

**Table 2** Characteristics of analyzed G1 nuclei and CTs.

**Table 3 Analysis of G1 nuclei and 45s rDNA.**

## Supplemental data

**Supplementary Figure 1** Differences in chromosome 9 arrangement (green) and 45s rDNA (yellow) activity in root and leaf. Models of individual arrangements were created based on raw data observation using BioRender.com.

**Supplementary Video 1** Rice root nucleus in G1 phase. Chromosome 2 (yellow) and chromosome 9 (green) were visualized using oligo-painting FISH. Nucleolus was stained by immunolabeling with fibrillarin (red). Nuclear DNA was counterstained with DAPI (blue)

**Supplementary Video 2** Rice leaf nucleus in G1 phase. Chromosome 2 (yellow) and chromosome 9 (green) were visualized using oligo-painting FISH. Nucleolus was stained by immunolabeling fibrillarin (red). Nuclear DNA was counterstained with DAPI (blue)

